# Viral etiology of Acute Respiratory Infections in Hospitalized Children in Novosibirsk City, Russia (2013 – 2017)

**DOI:** 10.1101/353037

**Authors:** Olga Kurskaya, Tatyana Ryabichenko, Natalya Leonova, Weifeng Shi, Hongtao Bi, Kirill Sharshov, Eugenia Kazachkova, Ivan Sobolev, Elena Prokopyeva, Tatyana Kartseva, Alexander Alekseev, Alexander Shestopalov

## Introduction

Acute respiratory infections (ARIs) pose a significant public health problem worldwide, causing considerable morbidity and mortality among people of all age groups [1]. Children are on average infected two to three times more frequently than adults. [2]. There are more than 200 respiratory viruses that can cause ARIs. Respiratory syncytial virus (RSV), human rhinovirus (HRV), human metapneumovirus (HMPV), human parainfluenza virus (PIV), human enterovirus (EV), influenza virus (IFV), human coronavirus (CoV), adenovirus (ADV), and human bocavirus (BoV) are the most common viral agents associated with ARIs, accounting for around 70 % of ARIs [3, 4]. The frequency of mixed respiratory viral detection varies from10%to 30% in hospitalized children [5–7]. In addition, several new human respiratory viruses have been described in recent years, including human metapneumovirus [8, 9], human bocavirus, and novel human coronaviruses, including severe acute respiratory syndrome coronavirus (SARS-CoV) [10], human coronaviruses NL63 (HCoV-NL63), HKU1 (HCoV-HKU1) [11], and Middle East respiratory syndrome coronavirus (MERS - CoV) [12].

Although the majority of ARIs are associated with respiratory viruses, antibiotics are often used in the clinical treatment of ARIs. As children with ARTIs often have similar clinical symptoms, studying the clinical characteristics of children with virus-related ARIs and the spectrum of respiratory viruses will facilitate the development of precise treatments for ARIs [13]. Rapid diagnosis is important not only for timely treatment starting but also for the detection of a beginning influenza epidemic and the avoidance of unnecessary antibiotic treatment [14, 15].

Western Siberia is of great importance in ecology and epidemiology of emerging diseases. This territory was involved in the circulation of A/H5N1 and A/H5N8 avian influenza viruses in 2005 – 2017 [16, 17]. These viruses were spread by wild birds’ migration. Western Siberia is a place of crossing of birds’ migratory flyways wintering in different regions of the world: Europe, Africa, Middle East, Central Asia, Hindustan, and South East Asia. Therefore, there is high probability of emergence of reassortant strains between human and animal influenza viruses, as well as emergence of local outbreaks of human morbidity caused by uncommon variants of influenza viruses. Furthermore, Novosibirsk is the largest transport hub in this part of Russia with numerous international connections, that is important for the spread of ARIs [18, 19].

The prevalence of respiratory viruses among children with ARIs differs in different regions and varies over time [20–24]. Thus, to better understand the epidemiology of Acute Respiratory Infections in Russia, we investigated etiology of ARIs in children admitted to Novosibirsk Children’s Municipal Clinical Hospital in 2013 - 2017.

## Materials and methods

### Ethics issues

All aspects of the study were approved by the Ethics Committee of the Federal State Budgetary Institution “Research Center of Clinical and Experimental Medicine” (2013-23). Accordingly, written informed consent was obtained from parents prior to sample taking.

### Patients and specimens

Participants enrolled to the study were children 0–15 years of age within 3 days of illness onset and had at least two of the following symptoms: fever, sore throat, cough, rhinorrhea, nasal congestion, sputum, shortness of breath, lung auscultation abnormalities, tachypnea, and chest pain. Paired nasal and throat swabs were collected from each patient admitted to Novosibirsk Children’s Municipal Clinical Hospital by hospital nurses. A total of 1560 samples collected during four epidemic seasons of 2013 – 2017 (October – April) were enrolled to the study. The epidemiological and clinical information including case history, symptoms, physical signs, and examination were included in a standardized questionnaire. The samples were placed immediately in viral transport medium (Eagle MEM, BSA and antibiotics) and stored at 4–8°C prior transportation to the laboratory (not more than 24 hours). Detection of respiratory viruses was performed immediately after delivery to the laboratory. All specimens were tested for 15 common respiratory viruses, including influenza virus types A, B (IFVA and IFVB), human parainfluenza virus (HPIV) types 1-4, human respiratory syncytial virus (HRSV), human metapneumovirus (HMPV), four human coronaviruses (HCo), human rhinovirus (HRV), human adenovirus (HAdV) and human bocavirus (HBoV), using a real-time PCR assay-kit.

### Nucleic acid extraction and reverse transcription

Viral nucleic acids were extracted from nasal and throat swabs using RNA/DNA extraction kit «RIBO-sorb» (Interlabservice, Russia) according to the manufacturer’s instructions. The extracted viral nucleic acid was immediately used to perform the reaction of reverse transcription using commercial kit “REVERTA-L” (Interlabservice, Russia).

### Virus detection

Detection of respiratory viruses, including HPIV 1-4, HRSV, HMPV, HCoV-OC43, HCoV-229E, HCoVNL63, HCoV-HKU1, HRV, HAdV, and HBoV was performed using a RT-PCR Kit «AmpliSens ARVI-screen-FL» (Interlabservice, Russia), and IFVA and IFVB virus detection was performed using a RT-PCR Kit «AmpliSens Influenza virus A/B-FL» (Interlabservice, Russia) according to the manufacturer’s instructions. Positive and negative controls were included in each run.

### Statistical analysis

Two-tailed chi-square test (two by two table) was performed to compare the infection rates for respiratory viruses among different age groups. P-value <0.05 was considered to be statistically significant.

## Results

### Patient characteristics

Totally, 1560 samples collected from patients with ARI during the period from December 2013 to April 2017 were enrolled in the investigation. There were 824 males (52.8%) and 736 females (47.2%), and the patient’s ages ranged from 3 months to 15 years. The majority of them (43.2%) were between 1 and 3 years old. The age distribution is shown in Table 1.

**Table 1.** Patient characteristics of 1560 children with ARI in Novosibirsk Children’s Municipal Clinical Hospital from 2013 to 2017.

| Characteristics of the population |  | ARI (%)* | Infected (%)** |  |  |
| --- | --- | --- | --- | --- | --- |
|  |  | Total<br>N= 1560 | Total<br>infection<br>N= 1128 | Single<br>infection<br>N= 965 | Co-infection<br>N= 163 |
| Gender | Male | 824 (52.8) | 601 (72.9) | 504 (61.2) | 97 (11.7) |
|  | Female | 736 (47.2) | 527 (71.6) | 461 (62.6) | 66 (9.0) |
| Age group<br>(years) | < 1 | 325 (20.8) | 237 (72.9) | 194 (59.7) | 43 (13.2) |
|  | 1 – 3 | 674 (43.2) | 523 (77.6) | 436 (64.7) | 87 (12.9) |
|  | 4 – 6 | 259 (16.6) | 201 (77.6) | 178 (68.7) | 23 (8.9) |
|  | ≥ 7 | 302 (19.4) | 167 (55.3) | 157 (52.0) | 10 (3.3) |

\*- Proportion of each group in all the samples
\*\* - Proportion of virus-positive samples in each gender or age group

### Detection of respiratory viruses

Among 1560 samples, 1128 (72.3 %) were found positive for at least one virus, and 432 (27.7%) were negative for all respiratory viruses tested (Table 1). There was no significant difference in the incidence of respiratory viral infection between boys (601/824; 72.9%) and girls (527/736; 71.6%) (χ^2^ = 0.345, p>0.05). The respiratory virus positive rate appeared to decrease with age. The lowest positive rate was observed in the age group of more than 6 years old (167/302; 55.3%), while the positive rates in age groups less than 6 years old were more than 70% (Table 1). Statistically significant difference was observed between the age group of more than 6 years old and other age groups (χ^2^ = 54.113, p<0.01). No statistically significant difference was observed among the age groups less than 1 year old, 1 – 3 years, and 4 – 6 years.

Among all the samples, single infections accounted for 61.9% (965/1560), while coinfections accounted for 10.4% (163/1560) with the lowest rate of incidence in children more than 6 years old compared to children younger than 6 years (χ^2^ = 20.389, p<0.01) (Fig 1).

**Fig 1.**
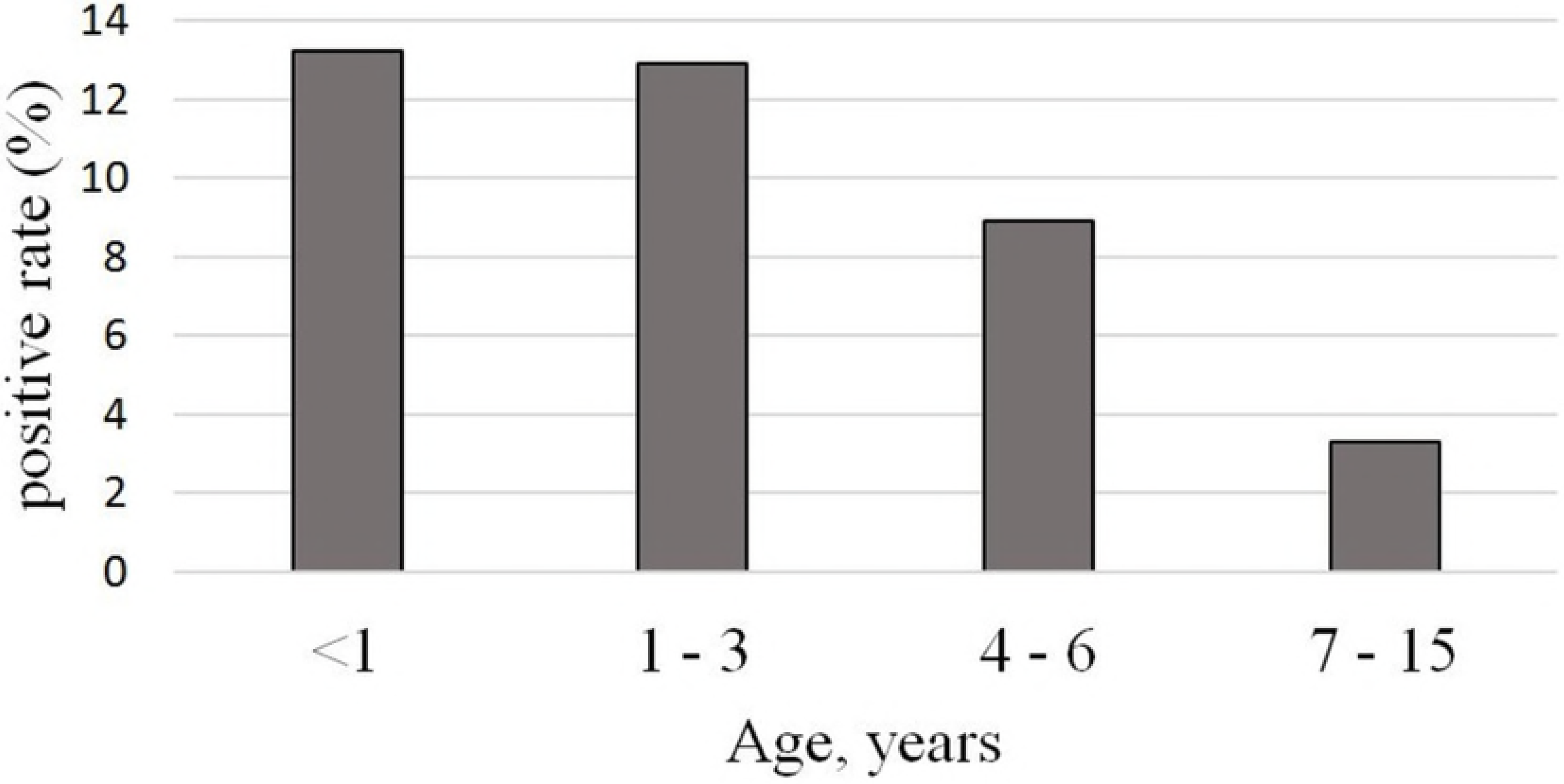
Viral co-infection rate in different age groups.

### Viral etiology

RSV and IFV were the most frequently detected viruses with high incidence of 23.0% (358/1560) and 22.1% (344/1560), respectively, among all patients with ARIs. HRV was detected in 15.1% (235/1560), followed by HMPV, HPIV and HBoV with the detection rates higher than 5.0%. The positivity rates of HCoV and HAdV were lower than 5.0% (Fig 2).

**Fig 2.**
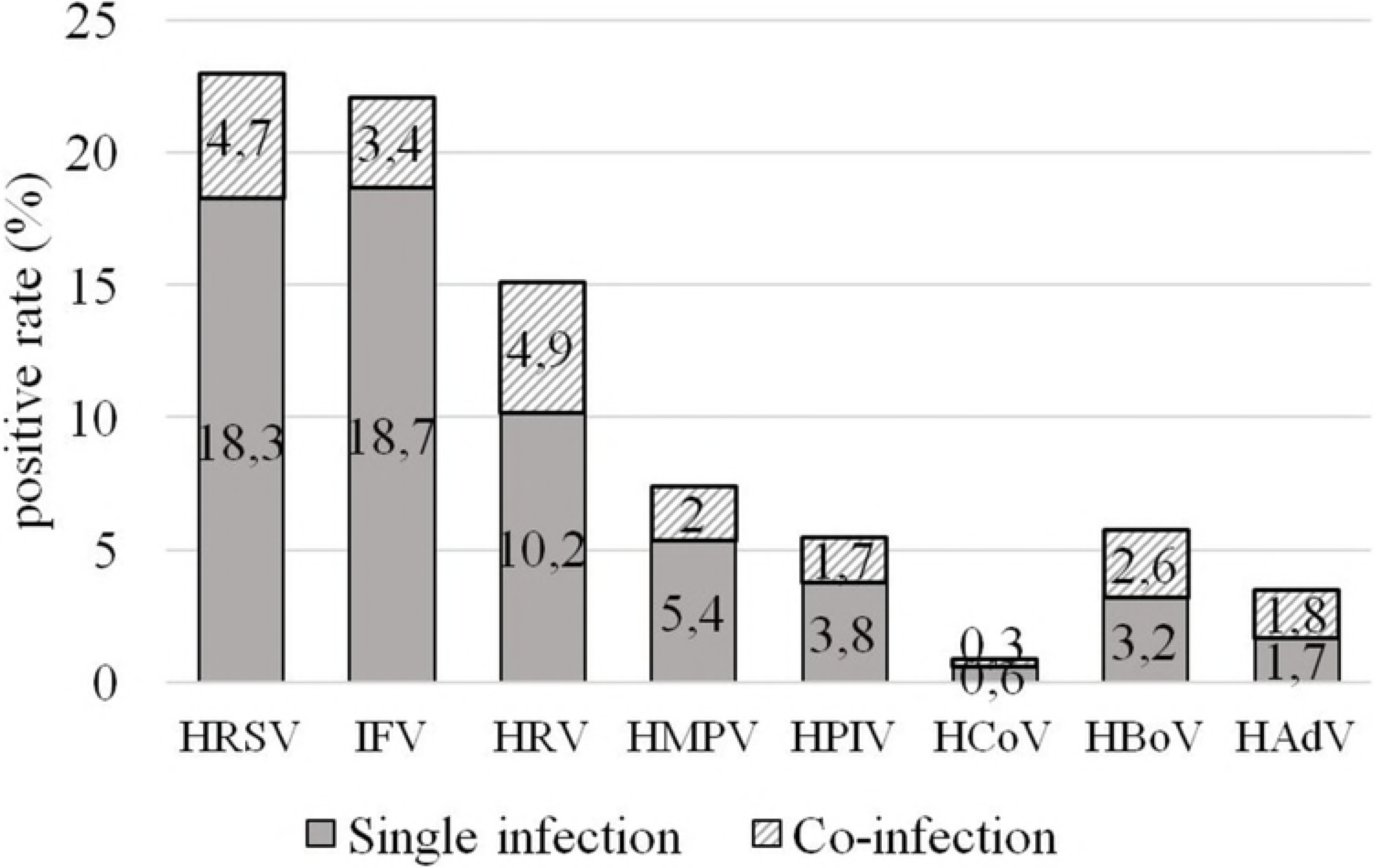
Detection rates of viral pathogens in single and co-infections in children with ARIs (2013 – 2017).

### Age and gender distribution

The data was analyzed with regard to the age and gender distribution of virus infection. Of the enrolled patients, 824 (52.8%) were male and 736 (47.2%) were female. All the patients were grouped into four age groups with different positive rate of viral infections. No difference in the etiological distribution of viral pathogens was observed between males and females.

Among detected respiratory viruses, HRV, HPIV, HCoV, and HAdV did not have statistically significant difference in the distribution among the different age groups. HMPV was detected in age group less than 1 year old much more frequently than in children older than 1 year (χ^2^ = 6.627, p<0.05). HBoV was significantly more frequently observed in children younger than 3 years old compare with children of 4 – 15 years old (χ^2^ = 28.523, p<0.005). The incidence of RSV decreased significantly with increasing age (p < 0.05) dropping from 35.1% in children younger than 1 year old to 5.3% in the school-age children (7 – 15 years old group), while the reverse relationship was observed for IFV (Fig 3).

**Fig 3.**
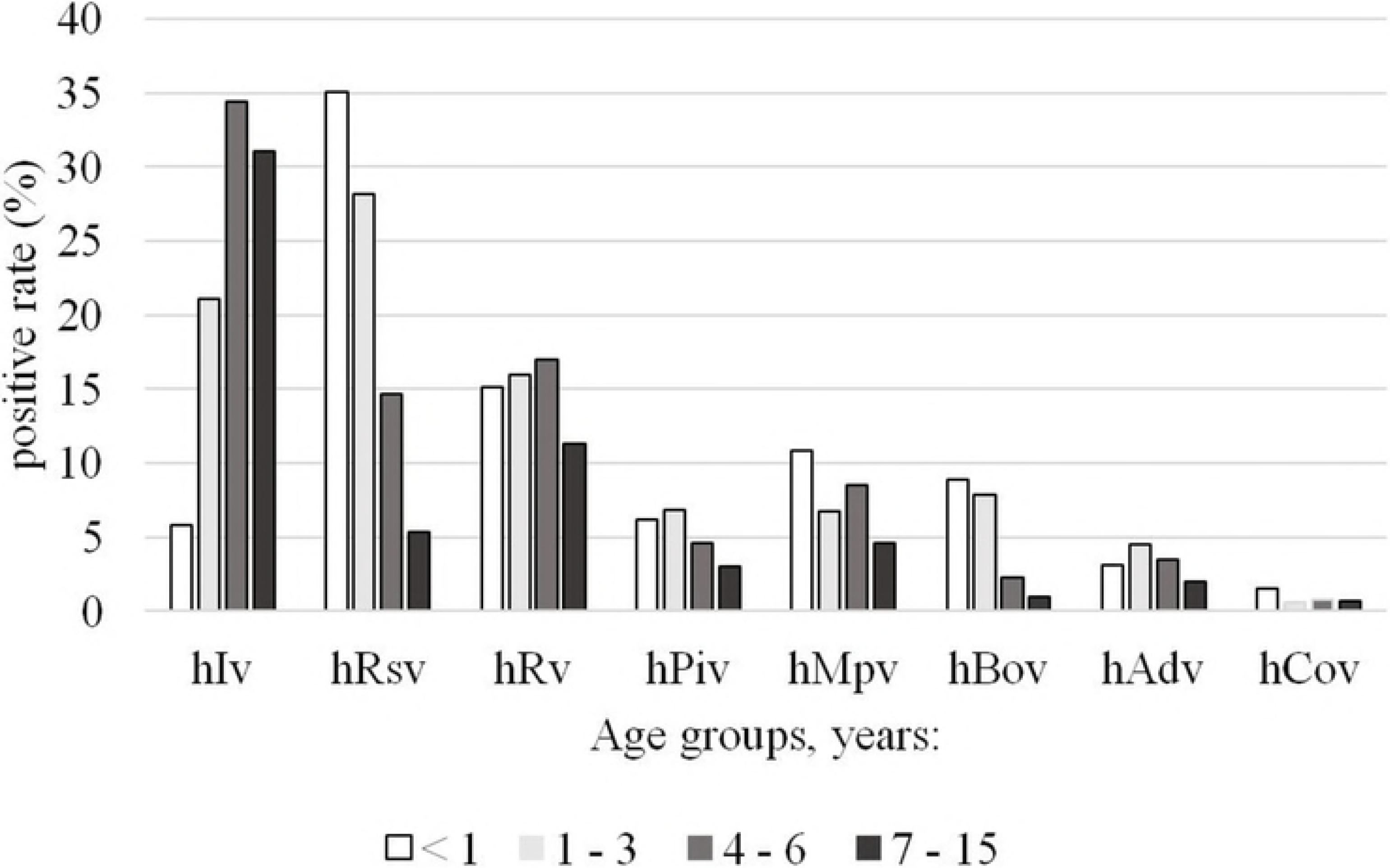
The distribution of respiratory viruses in different age groups.

### Seasonal distribution

The data was analyzed with regard to the seasonality. Figure 4 illustrates the monthly distribution of the most frequently detected viruses (HRSV, HRV and IFV) from 2013 to 2017. We have observed no considerable activity of Influenza viruses in 2013 – 2014 epidemic season, but increasing activity detected in subsequent years with peaks in February 2015, February 2016 and January and February 2017. HRSV exhibited marked peaks during each season: in January 2014, March 2015, December 2015 and March 2017. For HRV monthly distribution was relatively constant with only one clear peak in October – November 2014 (Fig 4).

**Fig 4.**
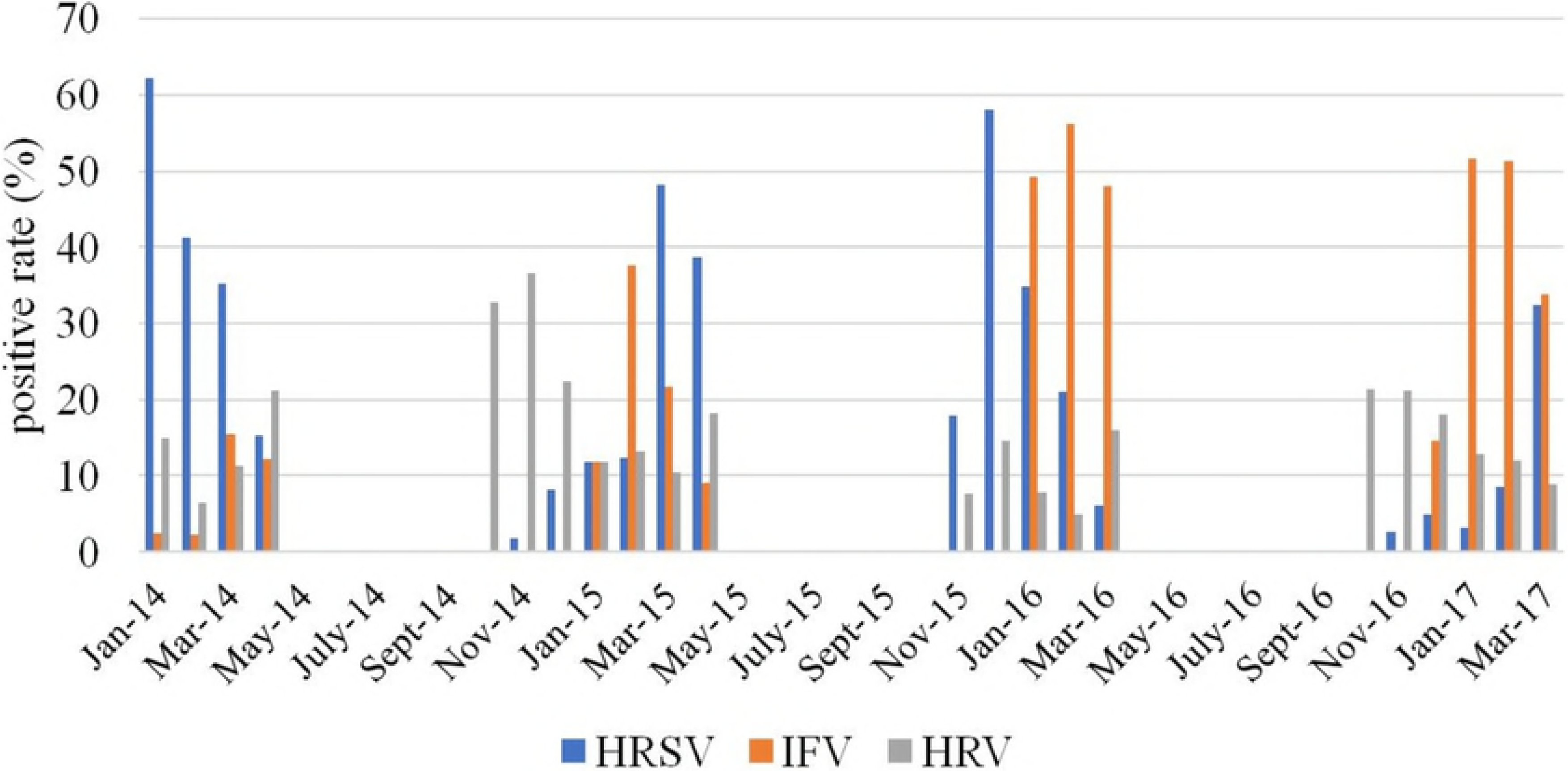
Monthly distribution of HRSV, HRV and IFV.

### Multiple Infections

Co-infections with two or more viruses were detected in 163 out of the 1128 (14.5%) positive samples (Table 1). Dual infections accounted for 11.4% (129/1128) of all positive samples and three viruses were detected in 3.1% (34/1128) of positive samples. Most co-infected patients were 0-6 years of age (12.2%, 153/1258) versus children older than 6 years (3.4%, 10/292). No significant difference was found for incidence of co-infections between the age group less than 1 year (13.2%, 43/325), 1 – 3 years (12.9%, 87/674), and 4 – 6 years (8.9%, 23/259). The most common combinations were HRSV/HRV, and IFV/HRSV, which amounted to 13.5% (22/168) and 12.3% (20/163) of all cases of co-infection respectively. Co-infection rate of each individual virus detected varied significantly. Viruses appearing most often in co-infections were DNA-viruses –HAdV and HBoV – in 52.7% (29/55) of cases of adenovirus detection and in 45.1% (41/91) of cases of bocavirus detection. HRV was the most often co-infected with HBoV (34.1%, 14/41) and HAdV (31%, 9/29). We have not detected any case of simultaneous infection of HAdV and HBoV. All occurring combinations of viruses are shown in table 2.

**Table 2.**
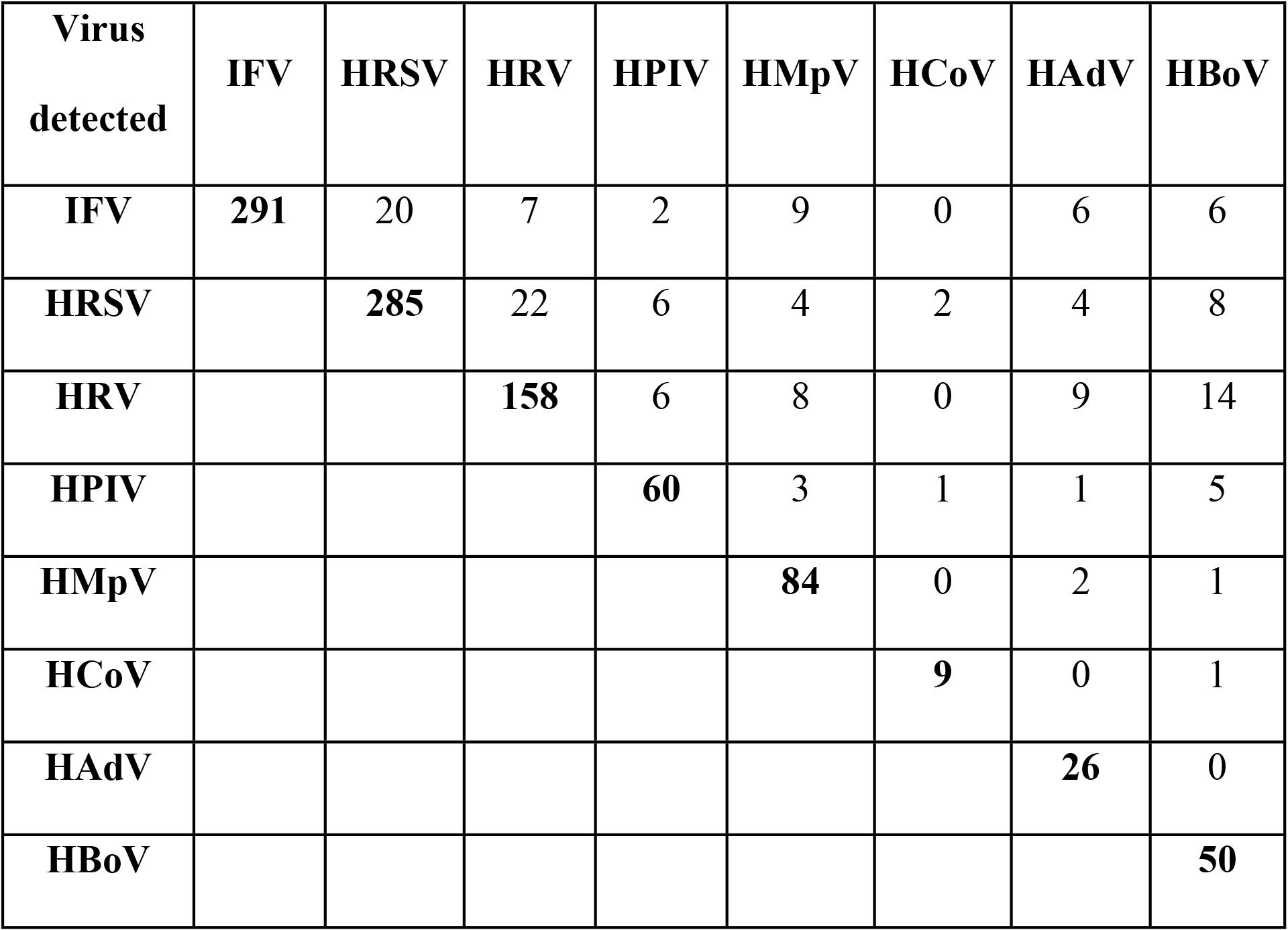
Detection of single and co-infection cases among 1560 children with ARI in Novosibirsk Municipal Clinical Hospital from 2013 to 2017.

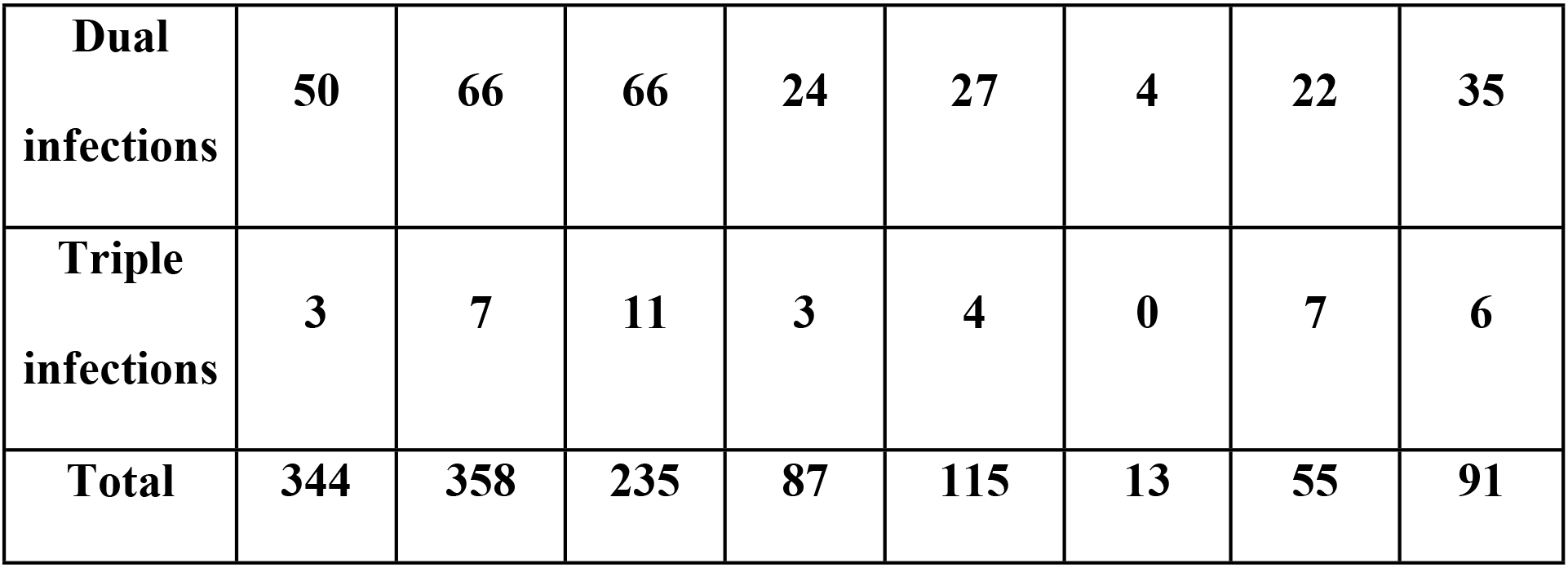

### Influenza viruses in etiology of ARIs

IFV was one of the most frequently detected viruses among children with ARIs with detection rate 22.1% (344/1560). The lowest detection rate of IFV was in the less than 1 year old age group (5.8%, 19/325). The incidence of IFV increased significantly with increasing patients’ age (p-value < 0.0001) showing 32.6% (183/561) in children older than 3 years.

During the study period the lowest influenza activity was investigated in 2013 – 2014 with positivity rate 6.9% of all positive samples. In 2014 – 2015 influenza virus detection was 17.2% while in 2015 – 2016 and 2016 – 2017 the detection rates were much higher and amounted to 32.2% and 30.2% respectively. Influenza A(H3N2) virus was predominant in 2013 - 2014 and 2014 – 2015 accounting for 57.9% and 69.8% of all influenza virus detections while in 2015 – 2016 82% of influenza viruses were A(H1N1) pdm09. We have not detected any influenza A(H3N2) viruses during 2015 – 2016 epidemic season. In 2016 – 2017 influenza type B detections (52%) predominating over type A (48%). Of influenza A viruses, all of them were A(H3N2) viruses (Fig 5).

**Fig 5.**
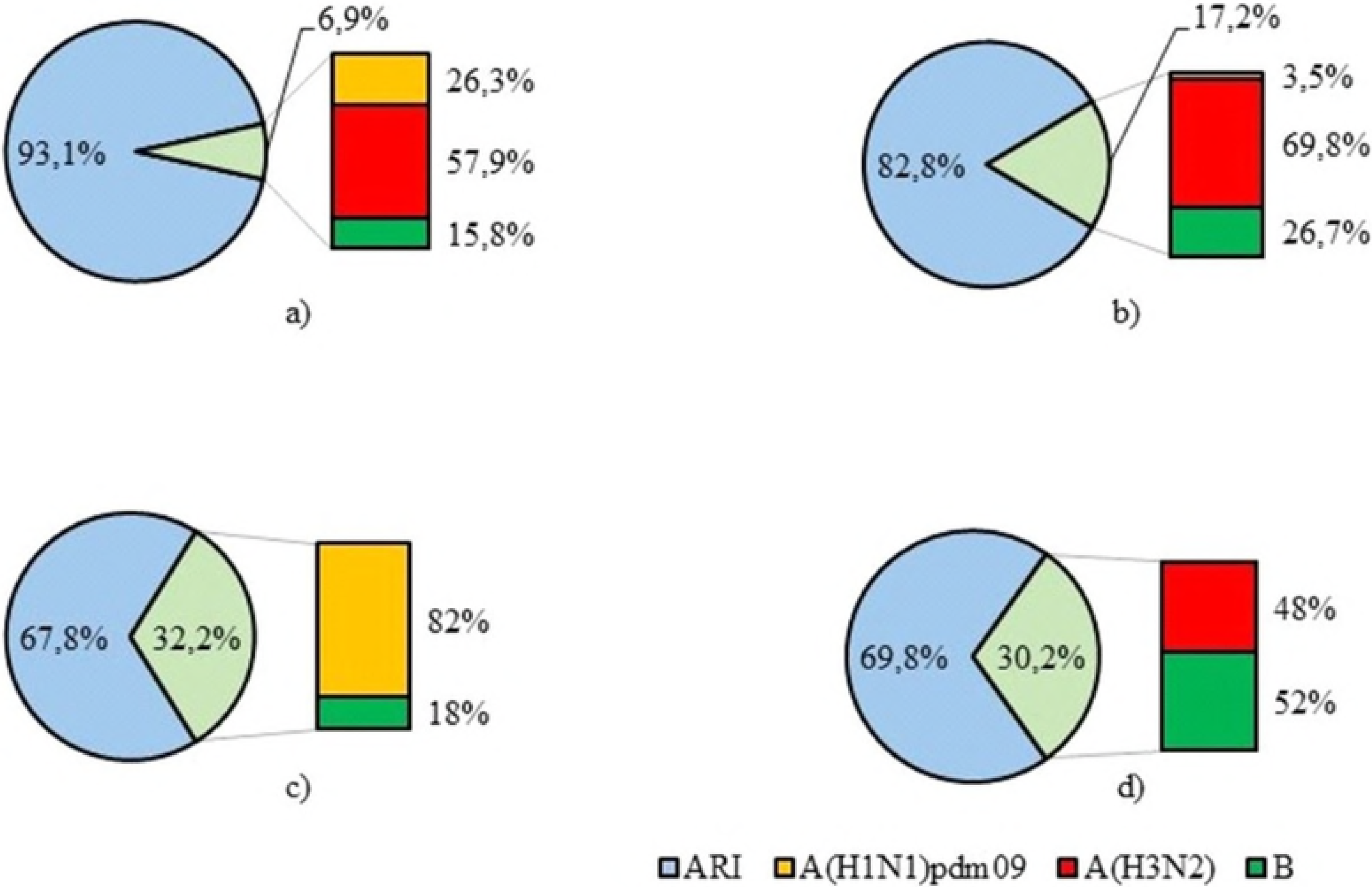
Distribution of influenza A and B viruses in 2013 – 2017.

## Discussion

Acute respiratory infections are a serious health and economic problem, causing high morbidity and significant economic losses due to temporary disability payments and medical costs. Children are the most susceptible group to the development of the disease. ARIs can lead to serious diseases such as bronchiolitis and pneumonia and sometimes even cause death in infants and children worldwide [25]. Nevertheless, most of the data on the epidemiological features and etiological structure of ARIs were from more-developed countries, and less is known about the etiology of ARIs in Russia. In the present study, we examined the viral etiology of ARIs in hospitalized children in Novosibirsk by Real-Time RT-PCR assay.

We detected at least one of the tested viruses in 72.3% (1128/1560) of the samples. In similar studies conducted in different regions of the world, the virus detection rate ranged from less than 50% to 75% [26–28]. For example, in studies performed in China, the proportion of positive samples in children with ARIs ranged from 37.6% to 78.7% [14,16,29,30]. The previous study of respiratory infections among children in European part of Russia revealed the 71.5% detection rate of respiratory viruses [31].

The percentage of the respiratory viruses’ detection varies in different years in different regions, which may be associated with climatic and environmental factors, population distribution, economic status and diagnostic methods used [1]. In addition, seasonality of sampling can also lead to differences in the level of viruses’ detection in various studies. Thus, Ju X. et al. carried out a study continuously from July 2011 through July 2013 and found 48.66 % of samples to be positive for at least one respiratory virus [32]. In contrast, we collected samples only during epidemic seasons of ARIs, so the positivity rate in our study was considerably higher. Furthermore, acute respiratory infections can be caused by viruses that are not yet known, as well as bacteria that have not been included in these studies [13].

We found that the prevalence of respiratory viruses did not differ between boys and girls (72.9% and 71.6% respectively), which confirms the absence of a gender-based susceptibility to respiratory viral infections [13]. However, we observed a decrease in the respiratory viruses’ detection rate with age, with the lowest detection rate in school-age children compared to children under 7 years of age (55.3% versus 76.4%, respectively). These data are consistent with findings obtained in other regions of Russia [31] and it may be due to decreased sensitivity to respiratory virus infections in older children.

Etiology of ARIs and the respiratory virus prevalence varies in different studies. In the United States, IFV, HRSV and HPIV were the most frequently detected [33]. Studies conducted in France have shown that metapneumovirus and respiratory syncytial virus are the most common [34]. IFV, HRSV and HRV were the most commonly detected respiratory viruses among children with ARIs in most regions of China [13]. The study of ARIs etiologic structure showed that HRSV, HRV, HPIV and IFV were registered significantly often among children in western part of Russia [31].

In our study the most common viruses detected were HRSV and IFV, followed by HRV. The age distribution of ARIs indicated that children under 3 years old were more likely to be infected by HRSV confirmed the importance of RSV in children with ARIs, especially in children < 4 years of age [35–38]. We observed a high rate of RSV-detections in 2013 – 2014 (44.4%), while in 2016 – 2017 it was less than 10% which could be due to the annual variation in the circulation pattern of RSV. Such year-to-year variation in the epidemiological patterns of viral infections confirms importance of the long-term study of the ARIs epidemiology [39].

Influenza virus is one of the major causative agents of respiratory disease in humans and leads to a more severe disease than the common cold which is caused by a different type of virus [40]. In temperate countries influenza outbreaks usually occur during the winter season. Finally, in our study, IFV (344/1560, 22.1%) was the second most frequent detected pathogen with markable seasonality in winter months. During the 2013-2014 epidemic season, influenza virus detection rate was low - 6.9% of all respiratory viruses, which was in accordance with the official influenza surveillance results of Ministry of Health, Russia [41]. In 2014-2015, influenza viruses were detected in 17.2% among all respiratory viruses. Herewith, the percentage of influenza A virus accounted for 73.3% and influenza B virus – 26.7% of all detected influenza viruses. At the same time, in Russia influenza B was the main etiological agent, accounting for 50.6% of all detected influenza viruses. In 2015-2016, influenza A(H1N1) pdm09 virus was predominant in Russia, accounting for 79% of all influenza-positive samples consistent with our results (82%). In 2016-2017 influenza A(H3N2) virus was dominant in Russia detected in 61.3% of all influenza virus cases [41]. In our study influenza A and B viruses were detected with approximately the same frequency (48% and 52% respectively).

With the introduction of molecular techniques, the detection of multiple co-infecting viruses has become common, though the prevalence of each virus varies between studies [42]. In our study detection rate of viral co-infection was 14.5% among the positive samples. According to the previous reports, the incidence of viral coinfection in children can reach 30% [43]. Most often co-infection found in children under the 5 years of age, that is associated with immaturity of the immune system and, thus, greater susceptibility to infection [44]. In our study, we significantly more frequently detected cases of simultaneous infection with two or more viruses in children under 7 years of age compared with children of school age (12.2% versus 3.4%), while there was no significant difference in the incidence of viral co-infection between the age groups of 0 – 1 year, 1-3 years and 4 – 6 years.

Thus, in our study we investigated the etiological structure of acute respiratory viral infections in hospitalized children in Novosibirsk, Russia, and evaluated age and seasonal distribution of the various respiratory viruses. Systematic monitoring of respiratory viruses is necessary to better understand the structure of respiratory infections. Such studies are important for the improvement and optimization of diagnostic tactics, as well as measures for the control and prevention of the respiratory viral infections.

## Acknowledgments

We thank the clinicians of Novosibirsk Children’s Municipal Clinical Hospital for their assistance with sample collection. We thank Dr. Galina Skosyreva and Elena Timofeeva (Department of propaedeutic of childhood diseases, Novosibirsk State Medical University, Novosibirsk, Russia) for their assistance with sample collection.

